# Burst Estimation through Atomic Decomposition (BEAD): A Toolbox to find Oscillatory Bursts in Brain Signals

**DOI:** 10.1101/2024.08.19.608642

**Authors:** Abhishek Anand, Chandra Murthy, Supratim Ray

## Abstract

Recent studies have shown that brain signals often show oscillatory bursts of short durations, which have been linked to various aspects of computation and behavior. Traditional methods often use direct spectral estimators to estimate the power of brain signals in spectral and temporal domains, from which bursts are identified. However, direct spectral estimators are known to be noisy, such that even stable oscillations may appear bursty. We have previously shown that the Matching Pursuit (MP) algorithm, which uses a large overcomplete dictionary of basis functions (called “atoms”) to decompose the signal directly in the time domain, partly addresses this concern and robustly finds long bursts in synthetic as well as real data. However, MP is a greedy algorithm that can give non-optimal solutions and requires a large-sized dictionary. To address these concerns, we extended two other algorithms – orthogonal MP (OMP) and OMP using Multiscale Adaptive Gabor Expansion (OMP-MAGE), to perform burst duration estimation. We also develop a novel algorithm, called OMP using Gabor Expansion with Atom Reassignment (OMP-GEAR). These algorithms overcome the limitations of MP and can work with a significantly smaller dictionary size. We find that, in synthetic data, OMP, OMP-MAGE and OMP-GEAR converge faster than MP. Also, OMP-MAGE and OMP-GEAR outperform both MP and OMP when the dictionary size is small. Finally, OMP-GEAR significantly outperforms OMP-MAGE when the bursts are overlapping. Importantly, the burst durations obtained using MP and OMP with a very large-sized dictionary are comparable to that obtained using OMP-MAGE with a much smaller-sized dictionary in real data obtained from two monkeys passively viewing static gratings which induced gamma bursts in the primary visual cortex. OMP-GEAR yields slightly smaller burst durations, but all the estimated burst durations are still significantly larger than the duration estimated using traditional methods. These results suggest that gamma bursts are longer than previously reported. Raw data from two monkeys, as well as codes for both traditional and new methods, are publicly available as part of this toolbox.

## 1 Introduction

Oscillations in different frequency bands such as beta (13-26 Hz) and gamma (30-80 Hz) in brain signals are linked to various brain functions [1]. For example, both actual and imagery movements are associated with beta desynchronization [2]. Different regions of the brain, such as the frontal cortex, multiple parts of the motor system, and basal ganglia exhibit changes in beta power when various motor functions are performed [3, 4, 5, 6, 7, 8]. Likewise, gamma rhythm can often be induced in visual areas (such as the primary visual cortex (V1)) by presenting specific visual stimuli such as gratings, and both the power and center frequency of the gamma bursts depend on the properties of the visual stimuli [9, 10, 11] as well as cognitive factors such as attention [12, 13, 14, 15, 16].

Gamma rhythm is hypothesized to provide a mechanism for communication across brain areas [17] or provide a temporal reference for the spiking activity that codes information relative to the rhythm [18], for which the rhythm should be sustained for behaviorally relevant timescales. However, several studies have shown that beta and gamma oscillations occur as bursts of small durations of the order of 100-200 ms [19, 20, 21, 22, 23]. Some studies have suggested that gamma bursts spontaneously arise with matched timing and frequency to facilitate information flow on a cycle-by-cycle basis [24]. In some cases, even sustained rhythms may appear bursty in noisy data [25].

Traditional methods for detecting bursts include spectral methods which use the time-frequency power spectrum of the signal computed using methods such as the Continuous Gabor Transform (CGT) [26, 20] or Wavelet Transform [23, 27]. In these methods, bursts are detected as epochs in the time-frequency plane where the power (in the gamma frequency range) exceeds a particular threshold. In addition, some constraints on phase consistency may be imposed [26]. Recently, a spectral estimator has been proposed that uses a set of wavelets (superlets) with increasingly constrained bandwidths that are geometrically combined in order to maintain good temporal resolution of single wavelets and gain frequency resolution in upper bands [28]. However, these spectral estimation-based methods have a fundamental limitation since, in many cases, the spectral estimator follows an exponential distribution [29, 30], due to which the estimated power is low most of the time, with occasional time points with high power. This causes even sustained oscillations to appear bursty [31]. Another class of methods uses filtering-based techniques, where the signal is bandpass filtered in the frequency range of interest and smoothed with a window of suitable length. Then, the burst onset and duration is estimated by finding the time points where the estimated amplitude exceeds a particular threshold [21]. Another method used the Hilbert transform to estimate the instantaneous power [22]. Although these methods do not involve direct spectral estimation, the power estimation and thresholding step is mathematically equivalent to spectral estimation, which makes power-dependent estimation fundamentally bursty in nature.

We had previously shown that in synthetic data with injected gamma bursts of known lengths, traditional methods failed to pick up long gamma bursts, especially when the gamma power was not very high [31]. We then proposed a method based on the Matching Pursuit (MP) algorithm, which finds bursts completely in the time domain and is able to pick up longer bursts in synthetic data. MP decomposes a signal into a linear combination of functions (called atoms) taken iteratively from a large, overcomplete dictionary and identifies bursts based on the parameters of the chosen atoms. We found that MP based burst estimation was much more robust to the choice of threshold. In real data recorded from two monkeys who passively viewed achromatic gratings, the median gamma burst lengths were about 300 ms (three times longer than traditional methods), although the mode gamma burst length was 100 ms, suggesting that gamma oscillations tend to occur in short bursts with occasional long bouts [31]. Similar results were obtained for the gamma oscillations induced by hue patches [32].

Although MP outperformed traditional methods, it has some limitations. While convergence of MP is guaranteed, it is asymptotic, which means that for a finite number of iterations, the error still may be large [33]. Further, the residue (difference between the original signal and the reconstructed signal from the selected atoms) at each iteration is only orthogonal to the previously selected atom and may be correlated with other selected atoms, which means that residue can still be represented in terms of previously selected atoms [33]. One way to address this issue is to make the residue orthogonal to all previously selected atoms, which is achieved using Orthogonal Matching pursuit (OMP). Convergence of OMP is of finite order and has been shown to provide better performance in high precision applications [34]. Both MP and OMP perform well if the dictionary has atoms that have a high correlation with the signal to be decomposed, and hence, performance is limited by dictionary size. But, when a large dictionary size is used, these algorithms are very slow. Recently, Canolty and Womelsdorf (2019) proposed a Multiscale Adaptive Gabor Expansion (MAGE) algorithm where a parameter reassignment step is used to refine the selected atom, which they used along with MP to estimate bursts [35].

Here, we present a toolbox called BEAD (Burst estimation using Atomic decomposition), which implements four algorithms: MP, OMP, OMP-MAGE, and a new method called OMP-GEAR which performs parameter reassignment using a different approach as compared to OMP-MAGE. In addition, this toolbox also implements the traditional methods discussed above. While MP [31] and MP-MAGE [35] have been implemented previously, OMP, OMP-MAGE and OMP-GEAR are implemented for the first time for gamma burst duration estimation. Methods to generate synthetic data which has gamma bursts of known durations are included. In addition, raw data from the two monkeys used in Chandran et al. (2018) [31] is also provided. This comprehensive toolbox will allow researchers to test various burst estimation methods on synthetic and real data and quantify their respective complexity-performance tradeoffs.

## 2 Results

The generation of synthetic data has been explained in detail in our previous study [31]. We injected multiple non-overlapping gamma bursts of known durations in spontaneous Local Field Potential (LFP) data while ensuring that the total power in the gamma range was similar to the gamma power observed in real data. When the gamma power was high, methods based on Hilbert Transform and Wavelet transform performed well, although CGT based method failed even in this condition (see [31, Figure 3C]). However, when the injected gamma bursts had low power, even Hilbert and Wavelet-based methods failed, and only the MP-based method was able to estimate the duration of long bursts properly (see [31, Figure 3I]).

Figure 1A shows the median of the estimated burst duration as a function of the injected burst lengths. All four atomic decomposition-based methods estimate the burst duration with high accuracy. For example, for an injected burst length of 300 ms, the estimated burst lengths were 306±3.8 ms, 306±5.7 ms, 306±3.7 ms and 308±2.8 ms using MP with 5 million atoms (MP-5M), OMP with 1.5 million atoms (OMP-1.5M), OMP-MAGE with 1.5 million atoms (OMP-MAGE-1.5M) and OMP-GEAR with 1.5 million atoms (OMP-GEAR-1.5M), respectively. In contrast, Hilbert based method performed poorly, with an estimated burst duration of 84±4.6ms.

**Figure 1:**
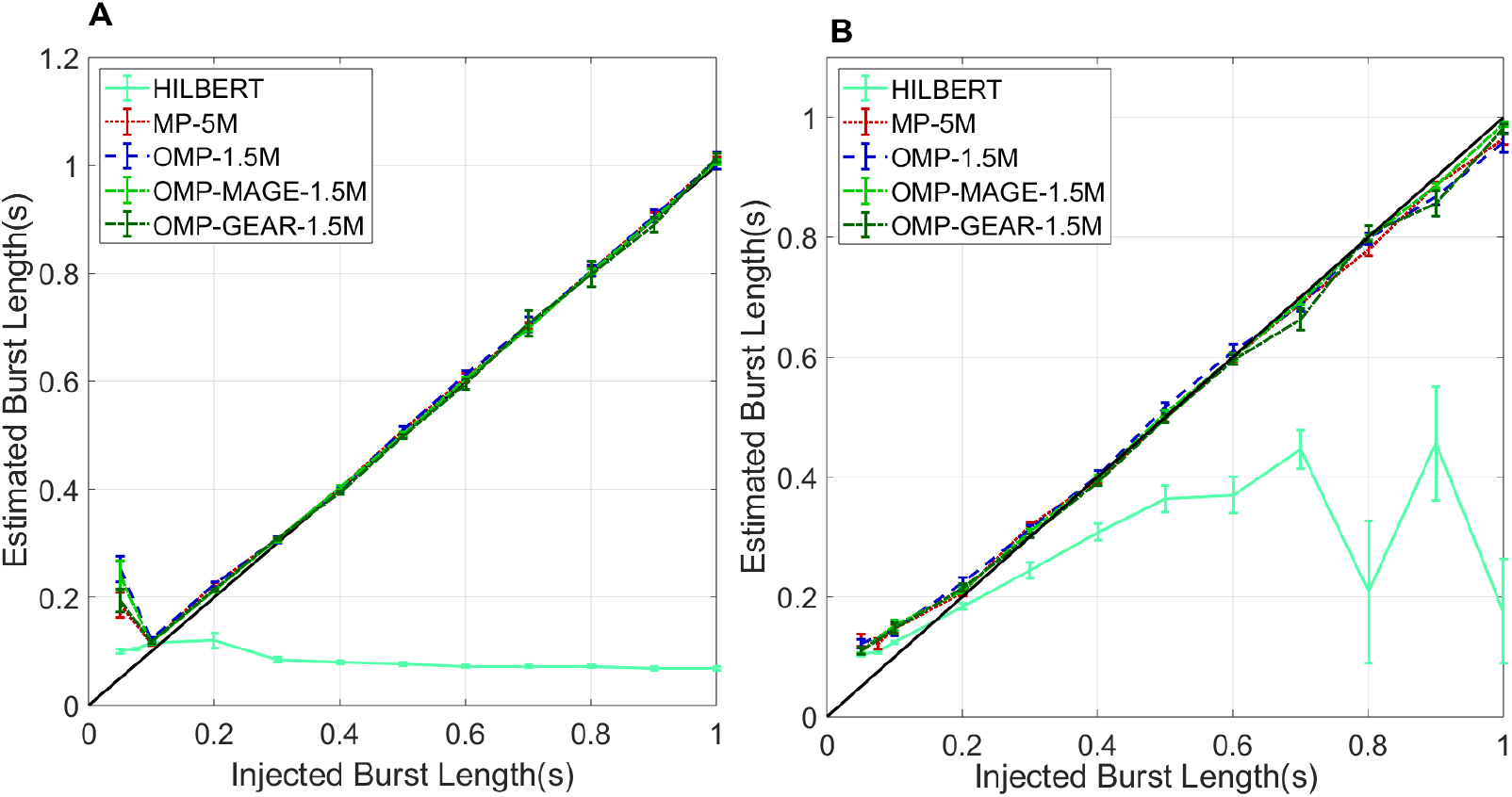
Comparison of gamma duration estimators. **A.** Injected vs. estimated burst lengths, when the injected burst has one-fourth the power of the real LFP in the gamma range. The bursts are non-overlapping, and the injected bursts are of duration 0.05 s,0.075 s and 0.1 to 1 s in steps of 0.1 s. The threshold fraction is set to 0.5. **B**. Injected burst vs. median burst lengths when the power of the injected burst matches with the real LFP in the gamma range. Overlap of bursts is allowed, and the injected bursts are of duration 0.05 s, 0.075 s, and 0.1 to 1 s in steps of 0.1 s. The threshold fraction is set to 0.5.

The only exception was when 50 ms bursts were injected. Because the total number of bursts was very high in this case (the number of bursts injected was increased in inverse proportion to the burst duration), two nearby bursts sometimes tended to get estimated as a single longer burst. Further, the sufficient conditions for sparse recovery are not met in this case (see Sec. 3.1), so it is not surprising that the atomic decomposition-based methods fail. However, the four atomic methods performed comparably in this condition, with estimated burst lengths of 185±22.8 ms, 252±24.2 ms, 240±28.1 ms, and 195±21.3 ms using MP with 5 million atoms (MP-5M), OMP with 1.5 million atoms (OMP-1.5M), OMP-MAGE with 1.5 million atoms (OMP-MAGE-1.5M), and OMP-GEAR with 1.5 million atoms (OMP-GEAR-1.5M), respectively. Although it appears that, in this case, the Hilbert method overestimated the burst length the least (at 100±3ms), the curve for Hilbert was almost horizontal, which means that it outputs the same burst duration for any injected burst duration.

To make burst estimation even more challenging, we removed the requirement that the bursts be non-overlapping. Figure 1B shows the results for this case. All four atomic methods continued to perform well, while the Hilbert-based method gave poor results.

In figure 1, OMP, OMP-MAGE and OMP-GEAR offer no improvement over MP, simply because we used a large dictionary (5 million atoms) and a large number of iterations, as determined in our previous study, to ensure that MP performed well for the synthetic data. However, such large dictionaries and iterations put severe constraints on the processing system and may not be desirable when memory or processing capacity needs to be minimized (for example, for real-time applications). Therefore, we tested the performance of the three atomic methods as a function of the number of iterations and dictionary size. For this, we simulated synthetic LFP data and injected bursts of 300 ms. Figure 2A shows the percentage reduction in the energy of the residue (the as-yet unexplained portion of the signal) as a function of the number of iterations. As expected, OMP, OMP-MAGE and OMP-GEAR converge much faster than MP, showing that if there is a constraint on the number of iterations, these methods can outperform MP. Figure 2B shows the estimated burst length as a function of the dictionary size. As the size of the dictionary reduces, OMP-MAGE and OMP-GEAR continue to perform well while MP and OMP significantly overestimate the burst duration. This illustrates the advantage of OMP-MAGE and OMP-GEAR relative to previous methods.

**Figure 2:**
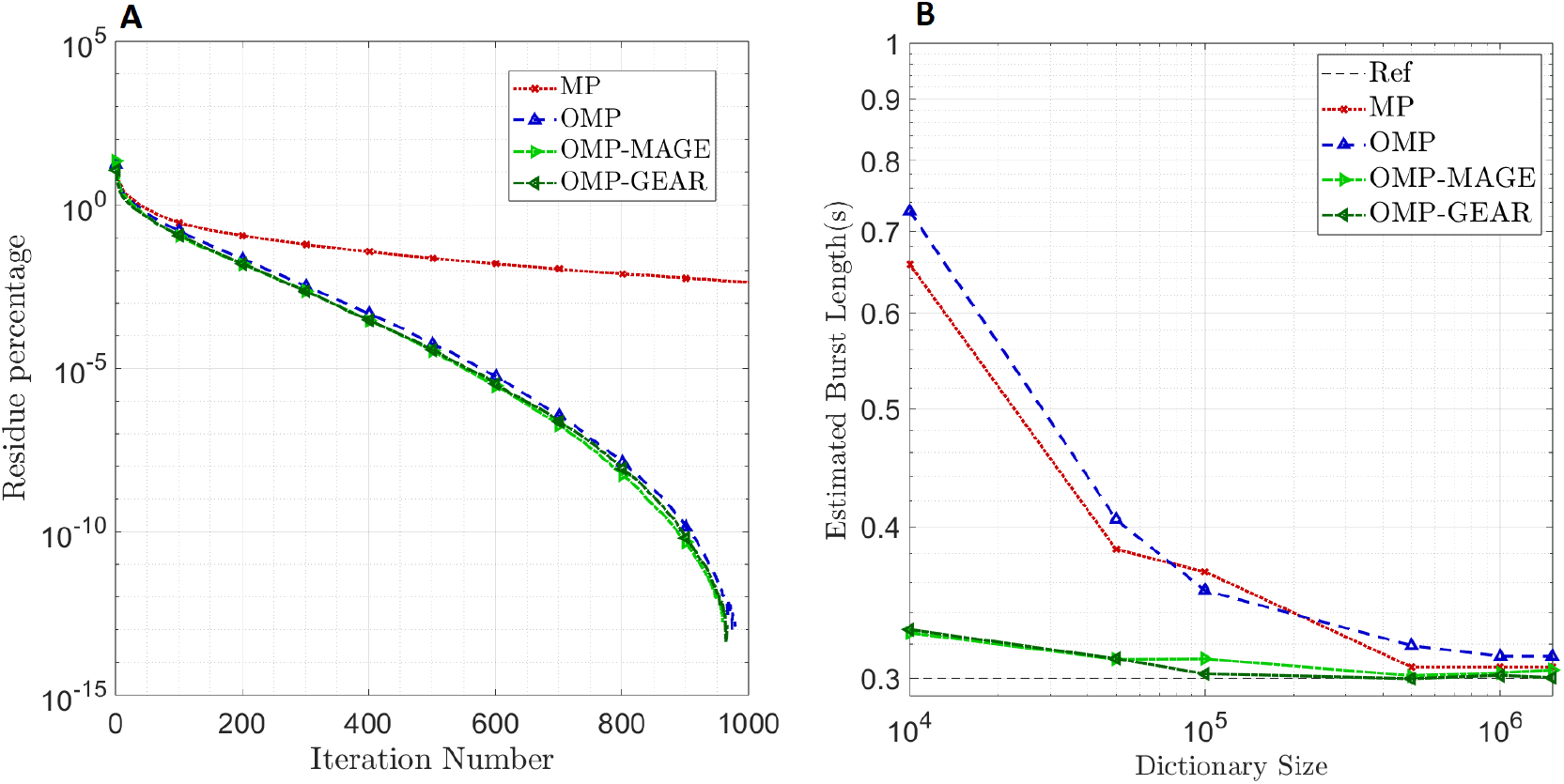
**A. Convergence analysis of methods:** Residue energy (percentage relative to the signal energy) vs. iteration number. **B. Performance of atomic methods as a function of the dictionary size:** Estimated burst durations for the three methods for various dictionary sizes when bursts of duration 300 ms are injected (shown as a dashed line with label Ref).

We tested the effect of increasing the extent of overlap between the bursts (see Fig. 3). For this, we increased the number of overlapping bursts of 300 ms duration. OMP-MAGE and OMP-GEAR were able to resolve multiple bursts better than MP and OMP. OMP-GEAR yielded lower error than OMP-MAGE when the number of bursts is high, showing that it is the best-performing algorithm.

**Figure 3:**
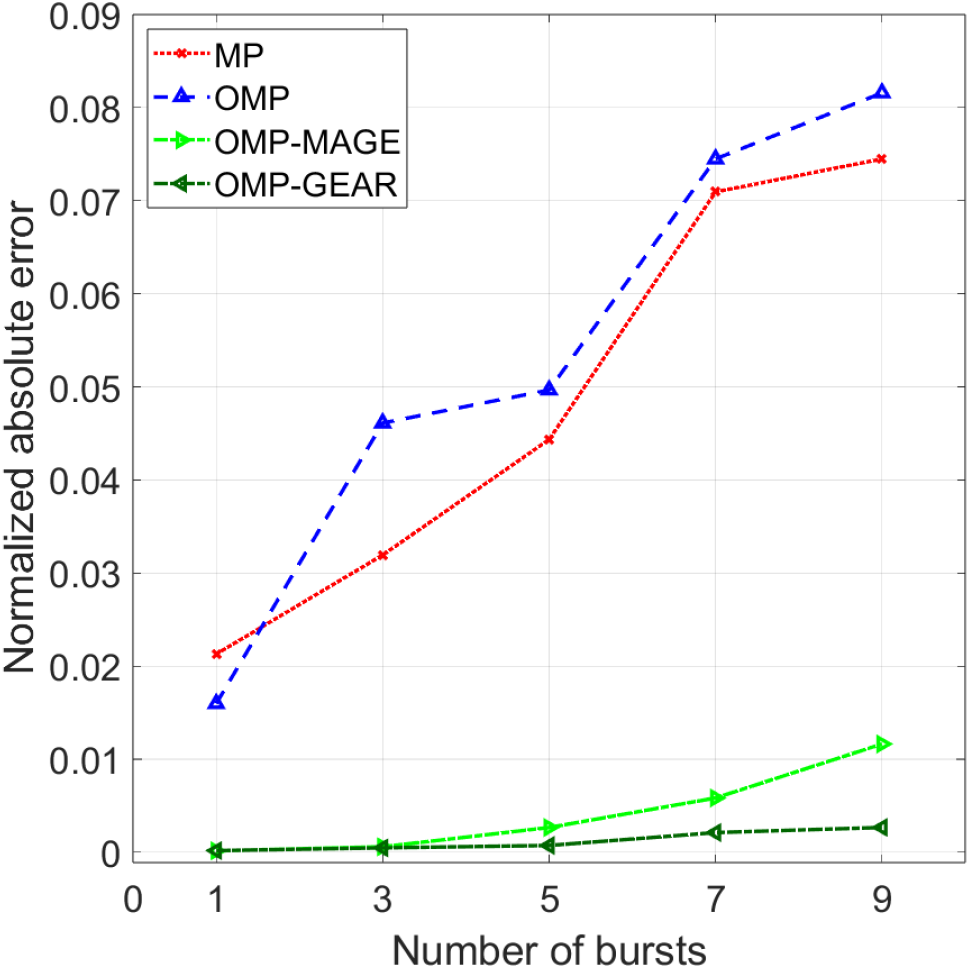
Performance of atomic methods as a function of number of bursts injected: Normalized absolute error for increasing number of bursts when bursts of duration 300 ms are injected.

Finally, we estimated the burst durations from real LFP data recorded from two monkeys. Results are presented (see Fig. 4) in the same format as [31, Figure 7]. MP, OMP and OMP-MAGE performed comparably, yielding median burst durations of 248.9±2.97 ms, 248.9±4.52 ms and 246.5±2.33 ms for monkey 1 and 287.2±9.97 ms, 300.0±4.85 ms and 302.9±7.85 ms for monkey 2, respectively. These durations were not significantly different from each other (p = 0.173, *χ*^2^=3.51 and p = 0.39, *χ*^2^=1.86 for monkey 1 and 2, Kruskal-Wallis test when MP, OMP, and OMP-MAGE were considered). OMP-GEAR yielded slightly smaller durations (227.5±1.27 ms for monkey 1 and 253.6±9.8 ms for monkey 2) compared to OMP-MAGE (p = 3.1 × 10^−6^, *χ*^2^ = 21.75 and p = 2.84 × 10^−6^, *χ*^2^ = 21.92 for monkey 1 and 2, pairwise Kruskal-Wallis test). The burst durations returned by the Hilbert-based method were much shorter (152±1.45 ms and 120±2.16 ms for monkey 1 and monkey 2, respectively).

**Figure 4:**
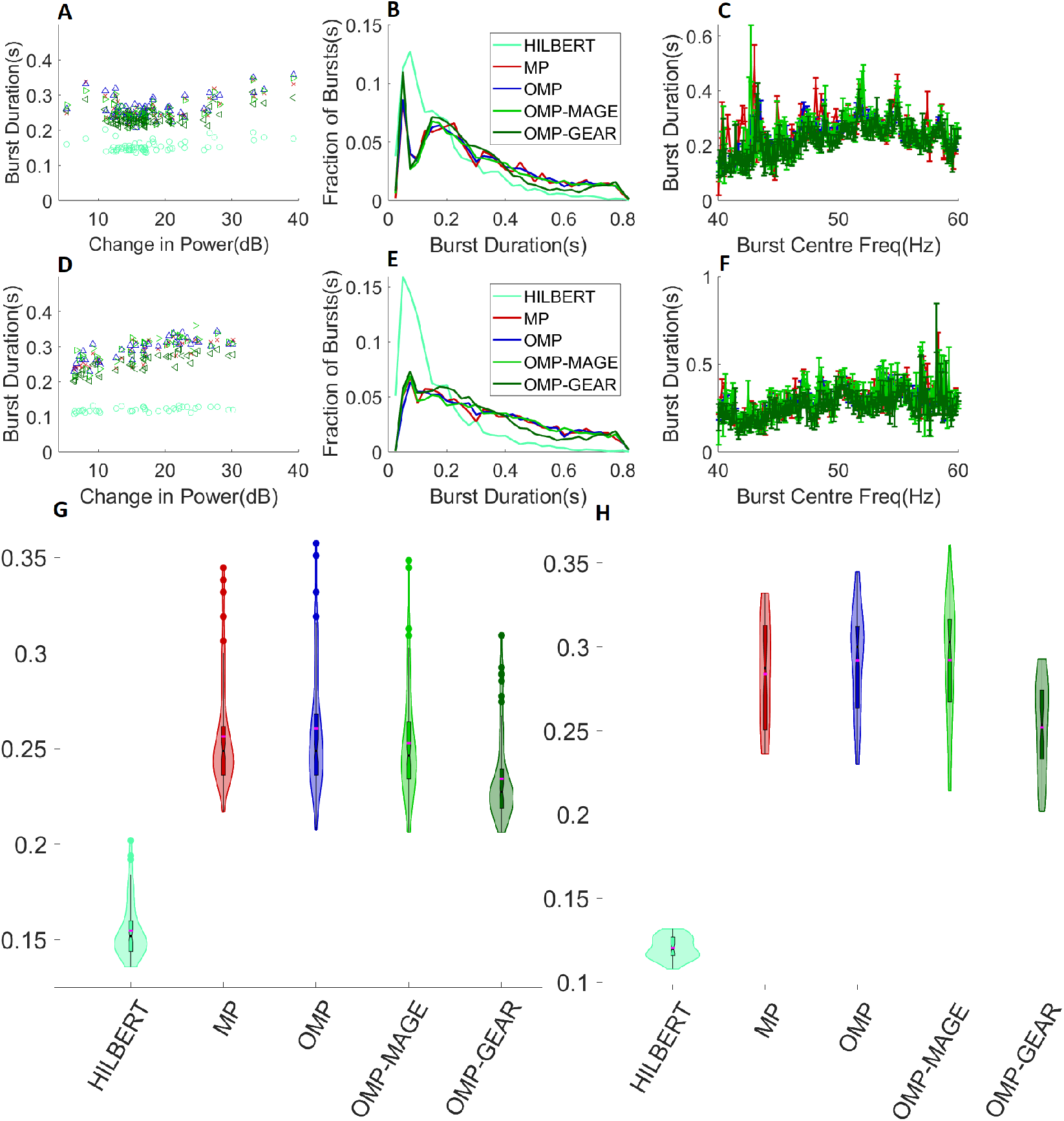
Gamma duration estimation from real LFP data. **A.** Median gamma duration for each electrode as a function of the change in the gamma power relative to spontaneous activity for monkey 1, for the four methods. **B**. Histogram of the gamma durations from all sites for monkey 1. **C**. Median gamma duration returned at each unique frequency by MP, OMP, OMP-MAGE, and OMP-GEAR, for monkey 1. **D**. Median gamma duration per site as a function of the change in power relative to the mean gamma power from spontaneous activity at that site for monkey 2. **E**. Histogram of gamma duration from all sites for monkey 2. **F**. Median gamma duration returned at each unique frequency by MP, OMP, OMP-MAGE, and OMP-GEAR, for monkey 2. **G**. Violin plots embedded with box plots of median burst lengths for all algorithms, for monkey 1. **H**. Violin plots embedded with box plots of median burst lengths for all algorithms, for monkey 2.

## 3 Discussion

We implemented three methods for burst duration estimation: OMP, OMP-MAGE, and OMP-GEAR. The OMP method is a modified version of the original method [34], where we incorporate a phase correction step to enable OMP to identify a suitable (orthogonal) pair of atoms along with the correct phase in each iteration. OMP-MAGE combines the MAGE approach [35] with OMP and incorporates the phase correction step mentioned above. Finally, we implemented a new method called OMP-GEAR. We found that when multiple bursts were presented, OMP-GEAR performed better than other methods in terms of correctly recovering the burst durations from synthetic data. All these methods gave longer median burst durations than traditional methods, validating the previous observation (using MP) that real data exhibits long burst durations [31]. The source codes and data is available at https://github.com/avianand8/GammaLengthProjectCodes.

### 3.1 Comparative Performance of MP, OMP, OMP-MAGE and OMP-GEAR

MP and OMP are greedy approaches for decomposing a signal as a linear combination of atoms from a dictionary. At each iteration, an atom is selected such that the energy in the residual signal decreases the most. The difference between MP and OMP lies in the least squares step in OMP, which makes the residue orthogonal to all the previously selected atoms. In turn, this results in the residual energy decreasing much faster in OMP compared to MP. There are other approaches for atomic decomposition such as basis pursuit [36], which pose atomic decomposition as a convex optimization problem.

We note that, in order to generate a dictionary of a given size, we first divide the parameter space into a fine dyadic grid, and generate atoms of the dictionary by sampling the grid points uniformly at random till the required number of unique atoms has been selected (see Sec. 4.3.1). The reason for using such a stochastic dictionary rather than a purely dyadic dictionary (which is generated by dividing the parameter space into a dyadic grid with the grid size such that the number of grid points equals the desired dictionary size, see Sec. 4.3.1) is because the latter introduces bias since the burst lengths are dyadic multiples of a base length. With OMP-MAGE and OMP-GEAR, we are not limited to the dictionary; the dictionary refinement step allows the algorithm to adjust the parameters of the atom to better explain the observation. Therefore, by carefully designing a fixed dictionary, OMP-MAGE and OMP-GEAR can yield better results even with small dictionary sizes.

From Fig. 1, we see that all four methods perform similarly for burst durations exceeding about 0.1 s in the non-overlapping bursts case, and for burst durations exceeding about 0.2 s in the overlapping bursts case. This is because the number of bursts injected is small relative to the size of the dictionary.^1^ Due to this, all sparse recovery-based methods perform nearly the same. For shorter burst durations (around 0.1 s), the number of injected bursts becomes large, causing the algorithms to slightly overestimate the burst duration, especially in the overlapping bursts case, since bursts in close proximity with similar phases could get merged as a single longer burst. At very low burst durations (around 50 ms), the number of bursts injected becomes too large (the mean number of bursts is 40) for sparse recovery techniques to work well, given the large size of the dictionary. A rule-of-thumb from the compressed sensing literature is that to recover an *N* -length, *k*-sparse vector using an *m*-dimensional linear observation obtained from noisy linear projections using an *m*×*N* measurement matrix, *m* = *O*(2*k* log *N*) is required. This condition fails when *N* = 1.5 × 10^6^, *k* = 40, since our measurements are *m* = 1024-dimensional vectors. Thus, all sparse recovery algorithms fail, as expected. Overall, this is an artifact of the way we have scaled the number of injected bursts as we reduce the injected burst length, but it helps to bring out the conditions under which the algorithms perform well and when they may fail.

Figure 2 shows that the orthogonal projections involved in the OMP-based algorithms allow them to pick sufficiently different atoms in subsequent iterations, resulting in a faster decay in the energy in the residue. Thus, OMP, OMP-MAGE and OMP-GEAR converge much faster than the original MP algorithm. Also, for a given dictionary size, OMP-MAGE and OMP-GEAR are not limited to the atoms in the dictionary. If the initially selected atom is sufficiently close to a true atom, OMP-MAGE and OMP-GEAR are able to refine the dictionary itself and replace the atom with one that is much closer to the true atom. Hence, as the dictionary size is reduced (consequently, the algorithm speeds up), OMP-MAGE and OMP-GEAR continue to return an estimated burst length that is close to the injected burst length (300 ms in this case), while both MP and OMP significantly overestimate the burst length. Additionally, from Fig. 4, we see that, on real data, MP performs comparably to the other three algorithms. This is because of the larger dictionary size (5 M) used with MP compared to the dictionary size (1.5 M) used with OMP and OMP-MAGE. OMP-GEAR yields slightly shorter burst durations, which could be because it is able to better resolve overlapping bursts, but these durations are still much longer than the ones obtained using Hilbert-based methods. Overall, atomic decomposition methods far outperform spectral methods such as the Hilbert transform-based method, showing that estimating the burst duration in the time domain itself is better than frequency domain based methods.

It must be noted that although atomic methods seemingly overestimate the burst durations occasionally, this is not necessarily incorrect, as there is ambiguity in the definition of a burst. If there are two nearby bursts that have a consistent phase and frequency relationship, they may be better represented by a single longer burst. Such a situation may arise physiologically if a sustained rhythm has an amplitude modulation, or if an underlying generator oscillates at a low frequency and generates bursts periodically (for example, see [25, Figure 1]). Indeed, the power of atomic methods is that they can potentially capture rich dynamics containing multiple bursts that are phase-aligned.

The regime where OMP-MAGE and OMP-GEAR outperform MP and OMP is when the number of injected bursts is moderately large (> 10) and the dictionary size is relatively small (about 1M). For much larger dictionary sizes (> 5M), the parameter reassignment used in OMP-MAGE and OMP-GEAR offers only a marginal advantage over OMP, as the dictionary already has sufficiently many atoms to represent the signal.

It is unclear why OMP-GEAR is able to better estimate the burst lengths than OMP-MAGE when multiple overlapping bursts are present. In both methods, we start with an initial estimate of an atom and subsequently refine it. OMP-MAGE uses a differentiation step (see Sec. 4.3.4) while OMP-GEAR (see Sec. 4.3.5) uses a different approach that skips the differentiation step. It is possible that skipping the differentiation step, which is prone to error or noise amplification, results in the improvement. In real data, we observe that the histograms in Figures 4B and 4E have a slightly lower fraction of bursts of durations around 500-600 ms and a slightly higher fraction around 200 ms compared to OMP-MAGE, suggesting that some bursts that were estimated as a single long burst by OMP-MAGE were instead resolved as two shorter bursts by OMP-GEAR. These observations lead us to believe that OMP-GEAR is a slightly better algorithm than OMP-MAGE, as it is able to resolve the bursts better.

### 3.2 Pros and Cons of Atomic Decomposition-based Methods

As seen previously, atomic decomposition-based methods can capture relationships between multiple bursts if they have some phase alignment. Also, depending on the signal characteristics, we can include other types of atoms in the dictionary such as chirp signals. Gamma oscillations are known to start with a high frequency on onset, which decays and stabilizes after some time. This transient behavior can potentially be better represented with chirps. Gamma oscillations may end more abruptly than the Gaussian window decay. One can include sinusoids multiplied by a rectangular window to capture these abrupt changes. The use of atomic decomposition and estimating the burst duration in the time domain provides the flexibility to include such atoms in the dictionary.

In terms of limitations, in our synthetic experiments, the injected bursts are Gabor waveforms and the dictionary that we used was also formed of Gabor atoms. The results could be different if injected bursts were generated using other waveforms but the dictionary was constructed using Gabor atoms. The greedy methods we have explored in this work have the limitation that if an incorrect atom is selected in a given step, the algorithms have no way of undoing the selection. Subsequent iterations that use a potentially incorrect residue may continue to pick incorrect atoms, cascading the error and resulting in a suboptimal signal representation. OMP-MAGE and OMP-GEAR alleviate this issue due to the dictionary refinement step, whereby these algorithms can correct for an incorrectly selected atom to some extent. However, it must be noted that even these algorithms require the initially selected atom to be sufficiently close to the true atom for the dictionary refinement to be effective. In turn, this means that the initial dictionary should sample the parameter space sufficiently finely to ensure that the dictionary contains an atom that is close to the true atoms. Finally, due to the large dictionary size requirement, these methods are computationally more expensive than traditional spectral decomposition-based methods.

## 4 Materials and Methods

### 4.1 Dataset Description

The dataset used in the study [31] consists of the Local Field Potential (LFP) collected from two adult female bonnet monkeys (macaca radiata), aged 13 and 17 years. The LFP signals were recorded by a 10 × 10 microelectrode array (96 active platinum electrodes; Utah array; Blackrock Microsystems) surgically implanted under general anesthesia. The Utah array was placed in each monkey’s primary visual cortex (area V1) of the right cerebral hemisphere (approximately 15 mm from the occipital ridge and 15 mm laterally from the midline). The receptive fields of neurons recorded from the microelectrodes were centered in the lower left quadrant of the visual field with eccentricities between 1.3 and 4.5 degrees.

The LFP signals were recorded while the monkeys performed a reward-based task where they fixated on a small spot in the center of the LED screen (BenQ XL2411, 1280 × 720 resolution, 100 Hz Refresh rate, linearized for contrast), while a large static grating of radius 4.8 degrees covering all the receptive fields was shown on screen. These grating were shown at five contrast levels: 0, 25, 50, 75, 100.

### 4.2 Synthetic LFP Model

Synthetic LFP is generated by injecting a random number of Gabor atoms [31] of the required duration (burst length) into the spontaneous LFP (denoted by *ϵ*). The model is given by

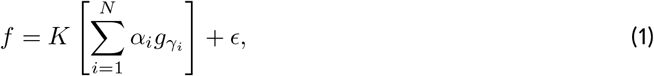

where 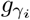 is a Gabor atom (the construction of 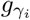 is explained in Sec. 4.3.1) with parameters *γ*_*i*_ = (*s*_*i*_, *u*_*i*_, *ξ*_*i*_). The number of injected bursts, denoted by *N*, is drawn from a Poisson distribution with a mean equal to the ratio of stimulus period duration to the burst length injected. Thus, the average number of bursts injected is inversely proportional to the burst length. The value of *u*_*i*_ is sampled from a uniform distribution over the range of the burst time. If any pair of bursts overlap in time, one of them is deleted. The value of *s*_*i*_ is set as 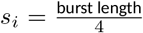. Note that, for Gabor atoms, *s*_*i*_ denotes the standard deviation of the Gaussian window, and the burst duration is well approximated by four times the standard deviation. Hence, this choice of *s*_*i*_ results in bursts of the desired length. The value of *ξ*_*i*_ is sampled from a uniform distribution in gamma range (40-60 Hz). The value of *α*_*i*_ is sampled from a normal distribution with a certain mean and standard deviation. The mean is set equal to the difference in magnitude of the average FFT of the stimulus and the standard deviation of the spontaneous LFP standard deviation and the standard deviation is set to 10 percent of the difference. The quantity *ϵ* denotes the spontaneous LFP signal and acts as additive noise. Finally, the burst signal 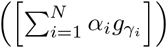 is scaled with a constant, *K*, such that the spectrum of the overall synthetic LFP matches that of the real LFP in the gamma range.

### 4.3 Burst Duration Estimation using Pursuit Algorithms

The procedure of burst estimation using pursuit algorithms attempt to solve the sparse approximation problem: 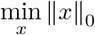 subject to *f* = *Dx*. This problem is NP-hard and is computationally infeasible even for moderate dictionary sizes. Pursuit algorithms such as MP and OMP solve this problem suboptimally, and is explained below. First, we explain the construction of the dictionary, *D*.

#### 4.3.1 Dictionary

All the sparse recovery algorithms require a dictionary to work. Before discussing the algorithms, we discuss how the dictionary is formed. Dictionaries constructed using orthogonal bases such as the Fourier basis provide a poor representation of functions well localized in time, while wavelet bases are not well suited to represent functions whose Fourier transform have narrow high-frequency support [33]. Therefore, dictionaries with waveforms well localized in both time and frequency are needed to represent nonstationary signals. Functions well localized in both time and frequency are called time-frequency atoms. A general family of time-frequency atoms can be generated by scaling, translating and modulating a single Gaussian window function *g*(*t*) ∈ *L*^2^(*R*) satisfying the constraint ∥*g*(*t*)∥ = 1. These time-frequency atoms can be parameterized by *γ* = (*s, u, ξ*), where *s, u, ξ* represent the scale, position, and frequency parameters, respectively. For a given *γ* ∈ Γ = *R*^+^ × *R*^2^, *g*_*γ*_(*t*) can be written as

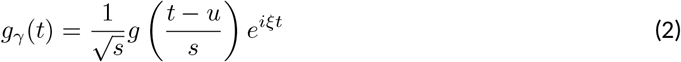

A dictionary containing a large number of atoms is constructed by gridding the parameter space Γ, and selecting an atom in that grid. We note that the parameter *γ* completely specifies the atom. From practical considerations, we limit the range of values of the scale, position, and frequency (*s, u, ξ*) parameters as *s* ∈ [1, *L*], *u* ∈ [0, *L*] and 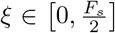, where *L* is the observation duration of the LFP signal *f* in seconds and *F*_*s*_ is the sampling frequency of *f*. The dictionary is constructed by dividing the parameter space Γ into a grid, and, if required, subsampling the grid to obtain the desired number of atoms.

##### Dyadic Dictionary

The most common approach is to form a grid in the dyadic scale, so that 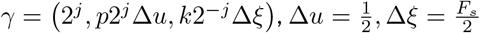, and *j, k* and *p* are integers spanning 0 *< j <* log_2_ *Z*, 0 ≤ *p < Z*2^*j*+1^ and 0 ≤ *k <* 2^*j*+1^, where *Z* = *LF*_*s*_. A dyadic dictionary has *O*(*Z* log *Z*) atoms [33].

##### Stochastic Dictionary

Using a dyadic dictionary introduces a bias because the scale parameter *s* is a power of 2 [37]. An alternative approach is to form a finely spaced uniform grid and sample the required number of atoms uniformly at random from this dictionary. This results in the so-called stochastic dictionary [37]. We have empirically observed that, on average, a stochastic dictionary produces better results than a dyadic dictionary.

#### 4.3.2 Matching Pursuit

Given a signal *f*, dictionary *D*, and sparsity level *M*, the MP algorithm [33] produces the following sparse approximation after *M* iterations:

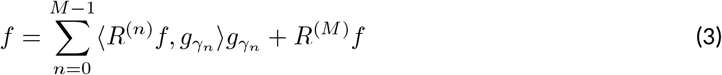

At each iteration, MP selects the atom 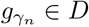 with maximum magnitude of correlation with the residue:

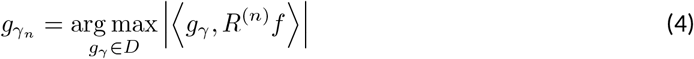

The residue in the first iteration is set as the signal itself. At each subsequent iteration, the residue is decomposed recursively: 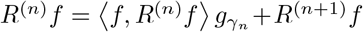. Due to (4), *R*^(*n*+1)^*f* is orthogonal to 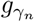 but not all previously selected atoms. Hence, the residue is correlated to the previous *n* − 1 atoms and can be represented in terms of these atoms also. Thus, an atom can be selected multiple times. As a consequence, MP does not make the best use of the selected atoms, and its convergence is only asymptotic.

#### 4.3.3 Orthogonal Matching Pursuit

OMP [34] is also an iterative algorithm that selects an atom at each iteration with the largest magnitude correlation with the residue. To obtain the best approximation of the signal with the selected atoms, at the *k*th iteration, OMP constructs a submatrix *D*_*k*_ of the original dictionary *D* formed using all the atoms selected till iteration *k*. The sparse vector *x*^*k*^ containing coefficients corresponding to the selected atoms is computed by solving the least-squares problem

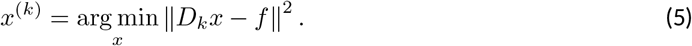

The solution to the above problem is 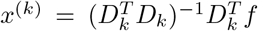, and the residue is updated as *R*^*M*^ *f* = *f* − *D*_*k*_*x*^(*k*)^. This makes the residue orthogonal to all the previously selected atoms, and we get the best approximation of the signal with the selected atoms. OMP is known to have a finite order of convergence [34]. It converges in a finite number of iterations; in fact, it converges in at most *m* iterations, where m is the length of the signal.

##### Phase correction

To properly account for the phase of the real-valued signal, we must form two dictionaries: one with the Gabor atom formed using the cosine function and the other with Gabor formed using the sine function. Consider a signal *f* = *g*(*t*) cos(*ξ*_*m*_*t* + *ϕ*) + *η*, where *η* is white gaussian noise, *m* = 1 or 2 are two potential frequencies. Suppose we have two dictionaries with two atoms each, Dcos = [*g*(*t*) cos(*ξ*_1_*t*), *g*(*t*) cos(*ξ*_2_*t*)] and Dsin = [*g*(*t*) sin(*ξ*_1_*t*), *g*(*t*) sin(*ξ*_2_*t*)]. Then, detecting the correct frequency is equivalent to non-coherent detection of frequency. It can be shown that when *ϕ* ∼ Unif[0, 2*π*], the minimum probability of error is achieved when we choose the frequency *ξ*_*m*_ such that

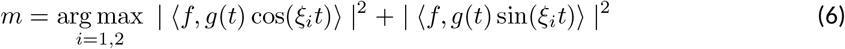

The phase of the signal is then estimated as

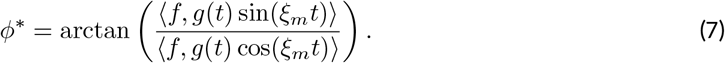

This phase estimate is the maximum likelihood estimate if the noise *η* is uncorrelated with the selected Gabor atom.

To adapt the OMP algorithm for real signals with unknown phase, we must thus start with two dictionaries containing the signals 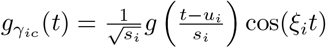 and 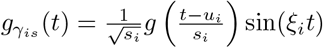. We normalize these functions to have unit norm while forming the dictionary. At iteration *k*, we select the parameter *γ*_*k*_ using the residue *R*^(*k*−1)^ from the previous iteration such that

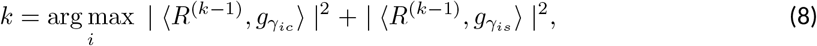

and the phase is estimated as

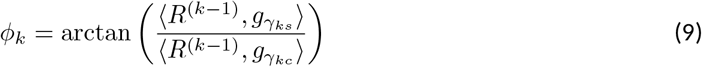

These two steps are necessary to represent signals using real-valued Gabor atoms.

#### 4.3.4 OMP-MAGE

MP and OMP select the atom with the largest correlation, but often the selected atom is not the true atom (depending upon the dictionary, it may be close to the true atom), as the dictionary does not contain all the possible atoms. Multiscale adaptive Gabor expansion (MAGE) [35] is a method in which the atom detected at each iteration is treated as an initial estimate. This estimate is further refined to move closer to the true atom. In MAGE, an atom is defined as

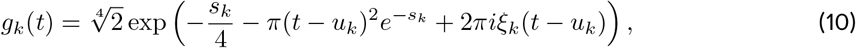

which is mathematically equivalent to (2). If we set *s*_*k*_ = 2 log_*e*_ *s*, the mapping is one-to-one, so for every *s*, there is a corresponding *s*_*k*_. The approach used in MAGE is one of curve fitting: the atom selected *g*_*p*_ is refined to a new atom *g*_*t*_ such that *g*_*t*_ fits the residue better at that iteration. Since *g*_*t*_ has three degrees of freedom *s, u* and *ξ*, we need three equations to update it. We explain the update equations below.

An analytical expression for the inner product between two atoms is given by

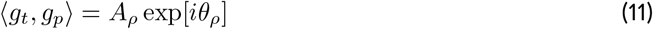

where

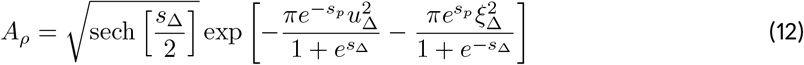

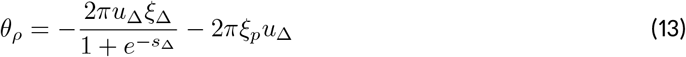

and *u*_Δ_ = *u*_*t*_ − *u*_*p*_, *ξ*_Δ_ = *ξ*_*t*_ − *ξ*_*p*_, *s*_Δ_ = *s*_*t*_ − *s*_*p*_. Using the chain rule, the derivative of the inner product with respect to a parameter *x*_*p*_ can be written as

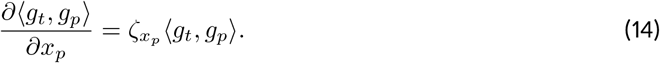

We can calculate 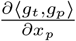 for all the three parameters *s*_*p*_, *u*_*p*_, *ξ*_*p*_ and ⟨*g*_*t*_, *g*_*p*_⟩ from the data. Also, we can obtain an analytical expression for 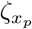 by differentiating (11) with respect to *s*_*p*_, *u*_*p*_, *ξ*_*p*_. By equating the analytical expression to the value calculated from the data, we arrive at the following closed-form analytical expressions for the parameters of the new atom:

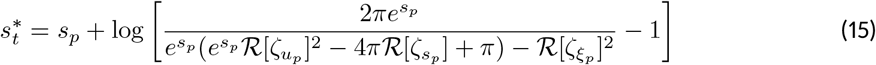

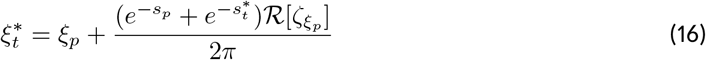

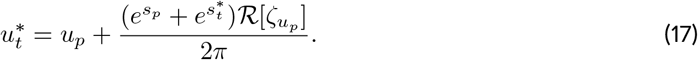

The above update equations are used as follows. At the *n*th iteration of the algorithm, suppose 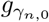 is the estimate of an atom that we get from MP or OMP using the residue computed at the start of that iteration, and suppose the true atom to be found (for example, the atom present in the signal that is closest to 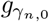) be 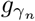. We can write the dot product between the residue at iteration *n* and the atom 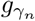 as

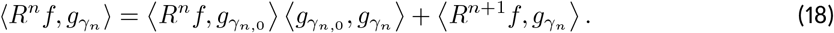

If 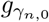 coincides with 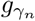, the interference term 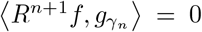. Using this, MAGE reassignment proceeds by neglecting the interference term and using (18) to reassign the atom parameters. This is further explained in Sec. 4.3.5.

##### OMP-MAGE vs MAGE

The original MAGE algorithm [35] was based on the use of MP to find the initial atom, whereas, in this work, we propose to use OMP to find the initial atom(s). Empirically, it is known that OMP has a better convergence rate than MP [34]. We also showed in Sec. 3 that OMP has much better convergence than MP. However, the use of MAGE in conjunction with OMP instead of MP requires us to form the dictionary differently. Specifically, we form one dictionary with the cosine function, 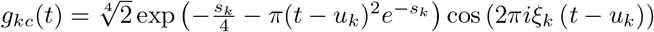 and the other with the sine function 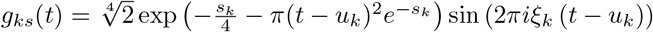 This allows us to use the real-valued update equations (15)–(17) to perform atom reassignment.

#### 4.3.5 Gabor expansion with atom reassignment (GEAR)

As discussed above, MAGE [35] finds a new atom *g*_*t*_ that is best matched with the residue at the current iteration by differentiating the inner product between the residue and the initial atom estimate rather than directly solving for the atom parameters. In effect, MAGE relies on a first-order Taylor series approximation of the atoms around the selected atom, and the resulting approximation error (see the note after (18)) can result in suboptimal parameter estimates. In this work, we present a novel alternative approach, where, instead of using a first-order Taylor approximation, we directly fit the residue to a new atom, *g*_*t*_, using the magnitude of the inner product between the residue and the initial atom, *g*_*p*_, found by OMP. The magnitude of the inner product between two Gabor atoms is given by

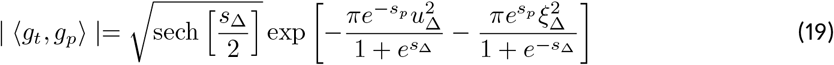

Suppose we want to fit the residue *R* at any iteration to a Gabor atom *g*_*t*_ and we have an initial estimate *g*_*p*_ with parameters (*s*_*p*_, *ξ*_*p*_, *u*_*p*_). We take another atom *g*_*p*1_ with parameters (*s*_*p*1_, *ξ*_*p*1_, *u*_*p*1_) where *u*_*p*1_ = *u*_*p*_ + Δ*u* and *ξ*_*p*1_ = *ξ*_*p*_ + Δ*ξ*. Since we wish to fit a true atom *g*_*t*_ to the residue, we take the ratio of the magnitude of the inner products of *R* with *g*_*p*_ and *g*_*p*1_, and simplify to get

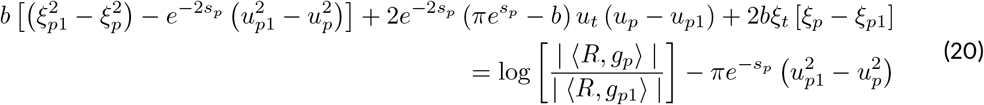

where 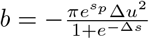. Defining intermediate variables 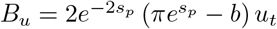 and *B*_*ξ*_ = 2*bξ*_*t*_, we get a linear equation in three variables *b, B*_*u*_, *B*_*ξ*_:

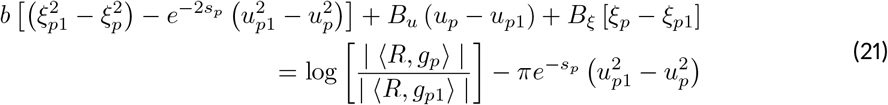

The right-hand side of (21) can be calculated from residue, initial estimate, and test atom *g*_*p*1_. Since the left-hand side has three unknowns, we need three equations. This can be obtained by taking two more testing atoms *g*_*p*−1_ with parameters (*s*_*p*_, *u*_*p*−1_, *ξ*_*p*−1_) and *g*_*p*1−1_ with parameters (*s*_*p*_, *u*_*p*1_, *ξ*_*p*−1_), *u*_*p*−1_ = *u*_*p*_ −Δ*u* and *ξ*_*p*−1_ = *ξ*_*p*_ − Δ*ξ*:

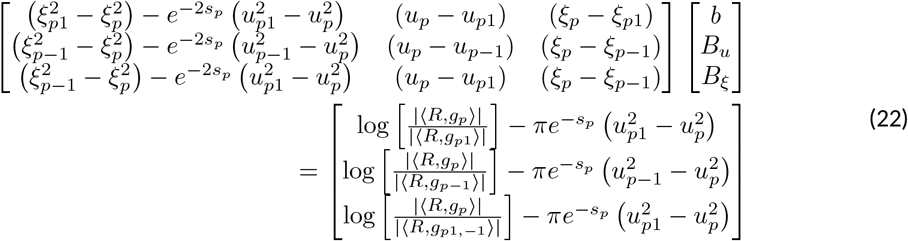

Using the system of equations above, we solve for the three variables *b, B*_*u*_, *B*_*ξ*_. Then, we can reassign the atom parameter values as

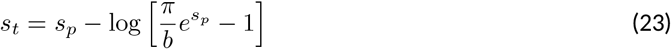

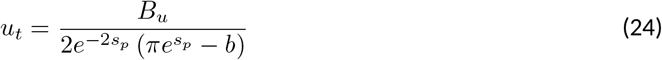

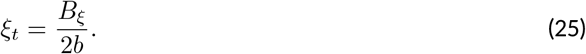

Note that, in order to obtain the system of equations in (22), we perturb only time and frequency parameters, i.e., we skip taking a finite difference with respect to the scale parameter. This is a pragmatic choice, as it allows us to obtain the above closed-form expressions for the atom reassignment. The parameters Δ*u* and Δ*ξ* must be chosen to be of a small value for the algorithm to converge; we use 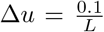 and 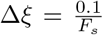, where *L* is the number of samples in the signal and *F*_*s*_ is the sampling rate. For signals with different parameters like sampling rate and number of samples, the parameters Δ*u* and Δ*ξ* can be tuned to get better results. We call this reassignment method used in conjunction with our implementation of OMP the OMP-GEAR algorithm.

From the sparse approximation of the signal using any of the three methods, we obtain a set of atoms, equivalently, a set of parameters that represent the signals. These parameters are used to estimate burst duration [31]. Atoms with frequency in the gamma range (40-60 Hz) and position parameters in the stimulus period (0-2 s) are selected as candidate gamma bursts. Finally, atoms whose coefficients exceed a threshold are included in the final estimate of the bursts. The threshold is set as a fraction of the mean of the largest coefficient obtained in the spontaneous period. The burst duration is set as four times the atom’s standard deviation, and the median of all these bursts across all trials is taken as the estimated burst duration. Burst durations exceeding 2 seconds are discarded as gamma bursts are typically more abrupt than the slow decay of a Gaussian window.

### 4.4 MP, OMP and OMP-MAGE Implementation

We use the implementation of MP by Piotr Durka et al. [37], which uses a stochastic dictionary instead of a dyadic dictionary as used in the original implementation by Mallat and Zhang [33]. This implementation is entirely written in C, with various optimizations to reduce computational time, making MP much faster than OMP and OMP-MAGE.

Our implementation of OMP, outlined above, has three major differences from conventional implementations of the algorithm. First, the (large) dictionary is not constructed in advance; a stochastic dictionary is formed on the fly using the dictionary size and sampling rate as inputs, inspired by [37]. Second, unlike conventional OMP where the inner product is computed with all atoms in the dictionary the atom with the highest magnitude correlation is selected, we form two Gabor dictionaries with the same parameters: one constructed using the cosine function and the other constructed using the sine function. Then we compute the inner product with all the atoms in the two dictionaries and choose the parameter that has the largest sum of squared of inner products of corresponding cosine and sine Gabor atoms. Third, we add a phase correction step by setting the phase of the sinusoid as the inverse tangent of the ratio of the inner products of the Gabor atom with sine and cosine functions. These enhancements are necessary for the correct representation of the signal.

### 4.5 Limitations of OMP and OMP-MAGE

The main limitation of OMP and OMP-MAGE is that these methods are slower than MP. There are two reasons for this: first, OMP and OMP-MAGE are both implemented in Matlab whereas MP is implemented in C, making the latter faster. Second, the MP implementation computes and stores inner products between dictionary atoms in advance, which speeds up the algorithm. However, in Matlab, the memory requirement for such a pre-computation is too large for practical implementation. To elaborate, in MP, atoms are computed when needed and only the atom parameters are stored, which requires multiple nested loops when implemented in Matlab, which makes the Matlab implementation of the same algorithm much slower than the C implementation.

There are faster ways to implement OMP, for example, using the inverse Cholesky factorization. When implemented in C, this can make our method much faster. As we saw earlier, OMP-MAGE offers comparable performance as OMP even with a much smaller sized dictionary (about 10x smaller). Hence, with a better implementation, the computational time of OMP-MAGE can be reduced, making it computationally superior to the MP-based method.

### 4.6 Burst Duration Estimation using the Hilbert Tranform

The analytical signal is formed from the given signal *f* as

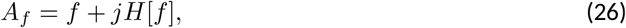

where *H* denotes the Hilbert Transform operator. This signal is band pass filtered in the gamma frequency range (40-60 Hz), and the envelope is found by taking |*A*_*f*_| ^2^. A burst is registered when the power exceeds three times the median power. The start and end of the burst are taken as the times when |*A*_*f*_| ^2^ falls below 1.5 times its median power.

## Footnotes

1 The number of bursts is a Poisson distributed random variable whose mean equals the stimulus period (2 s in our case) divided by the burst duration.

